# Vimentin networks at high strains

**DOI:** 10.1101/2025.10.06.680657

**Authors:** Shanay Zafari, Sophia Schirra, Raffaele Mendozza, Sascha Lambert, Kristian A. T. Pajanonot, Pallavi Kumari, Peter Sollich, Sarah Köster

**Affiliations:** Institute for X-Ray Physics, University of Göttingen, Göttingen, Germany; Institute for Theoretical Physics, University of Göttingen, Göttingen, Germany; Institute for the Dynamics of Complex Systems, University of Göttingen, Göttingen, Germany; Department of Mathematics, King’s College London, Strand, London WC2R 2LS, United Kingdom; Max Planck School “Matter to Life”, University of Göttingen, Göttingen, Germany

## Abstract

The cytoskeleton is crucial in maintaining cell shape and structural integrity. It consists of three types of filaments, including actin filaments and intermediate filaments (IFs) that exhibit distinctly different force-strain behavior: while IFs show nonlinear behavior with exceptional extensibility and remarkable resistance against rupture at high strains, actin filaments break at low strains. Here we address the question of whether the intriguing mechanical behavior of vimentin IFs translates to the network scale. We apply high strains to in vitro reconstituted networks using optical tweezers and find contrastive behavior for the two cytoskeletal networks: vimentin networks show strain stiffening behavior and respond in an elastic, solid-like manner with high forces opposing the active movement of the beads, whereas actin networks strain soften and fluidize at low forces. Our work highlights the complementary nature of the components of the cytoskeleton, which – in the cell – are partly co-localized and believed to constitute a composite biological material.

---

Biological cells are exposed to forces and deformations^1^ and their mechanical response to such impact is determined by the cytoskeleton, a complex biopolymer network of three filamentous components – microtubules, actin filaments and intermediate filaments (IFs). Among them, IFs are expressed in a cell-type specific manner and the most abundant representative of this protein family is vimentin, which is found in cells of mesenchymal origin. Each of the cytoskeletal filaments displays very specific mechanical properties. ^2–4^ In particular, IFs are very extensible, up to multiple times their original length,^5–7^ whereas actin filaments break at low strains of about 5% and forces of around 20 pN^8^ (see Supplemental Material (SM), Fig. S1 for data of stretched actin filaments). Furthermore, the force-strain behavior of vimentin IFs is highly non-linear and dependent on the loading rate: the filaments extend in a linear manner up to strains of 10–20 %. When pulled slowly, they then enter a plateau-like regime, where only little force is needed to extend them further, before they stiffen again. At faster extension rates, this plateau is almost completely missing and the filaments stiffen early on.^7^

In the context of cell mechanics the question arises of whether and how the filament properties translate to the network level. It was shown that networks of actin or microtubules break at low strain or stress, respectively, whereas vimentin networks remain undamaged. ^2^ While such bulk rheology studies typically probe the sample as a whole, optical tweezers-based active microrheology (AMR) is suited to strongly strain the networks locally, thus potentially reaching the non-linear regime of vimentin IFs. Using this approach, in entangled actin networks, it has been shown that above a critical strain rate of 3 s^−1^ and a critical protein concentration of 0.4 g/L,^9,10^ the networks behave in a non-linear manner, stiffen at short time scales, and then fluidize to a viscous solution. An interesting recent study compared the stress-strain response of actin and vimentin networks to that of composite networks.^11^ Both the elasticity and the viscosity of the composite networks were reported to be reduced compared to pure networks, contrary to what would have been expected from previous studies. The authors also observed emergent effects in the mixed networks during the initial response, whereas on long time scales the stress response was governed by actin.

Beyond in vitro reconstituted protein networks, AMR with optical tweezers has been applied to compare vimentin knockout (vim-/-) cells, which lack vimentin, and “ghost cells” where all components but the vimentin IFs have been dissolved.^12^ The authors find that upon straining the network by moving a bead within the cells, the vim−/− cells strain soften, whereas the ghost cells strain stiffen. Overall, the results from stretched single filaments,^7^ bulk rheology of in vitro networks^2,13^ and AMR in whole biological cells^12^ agree with each other (but contrast with Ref.^11^) and show that vimentin networks – and possibly IF networks in general – behave mechanically very differently to actin networks. Here, we bridge the gap between single filament mechanics and bulk shear rheology by AMR: we use optical tweezers to deform vimentin networks to large strains and find that they do indeed resist high forces, clearly exceeding what has been found for actin.

We embed passivated polystyrene beads in actin and vimentin networks and use optical tweezers to move the beads relative to the networks. The force acting on the bead is recorded and informs us about the networks’ response to the strain. In the experimental realization, the position of the trap is kept fixed and the stage is moved by a specified maximum displacement *s*_max_ at a specified constant speed 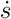 (see sketch and plot in Fig. 1(a) and (b, gray), respectively), whereby the bead imposes stress on the network and the displacement of the bead relative to the center of the trap is used to determine the force [Fig. 1(b, red)] via the known trap stiffness (see SM, Sec. I for a detailed description of the methods). After the extension phase, the position of the nanostage is kept fixed, and the bead is still trapped to investigate the relaxation of the network. We employ two different experimental protocols: we keep the total time for the extension phase constant at 1 s and vary the velocity of the stage 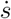 in proportion to the maximum displacement *s*_max_ of the stage; (2) we keep the maximum displacement constant and adjust the total extension time according to the chosen velocity. We choose probe particles that are about 4.5 times larger in diameter *a* = 2 *µ*m than the theoretically calculated mesh size *ξ* = 450 nm [Fig. 1(c)].^14,15^ Thus, the beads are well-captured within the mesh,^16,17^ while the required laser power for the AMR experiment remains in a reasonable range so as not to destroy the protein.

**Figure 1:**
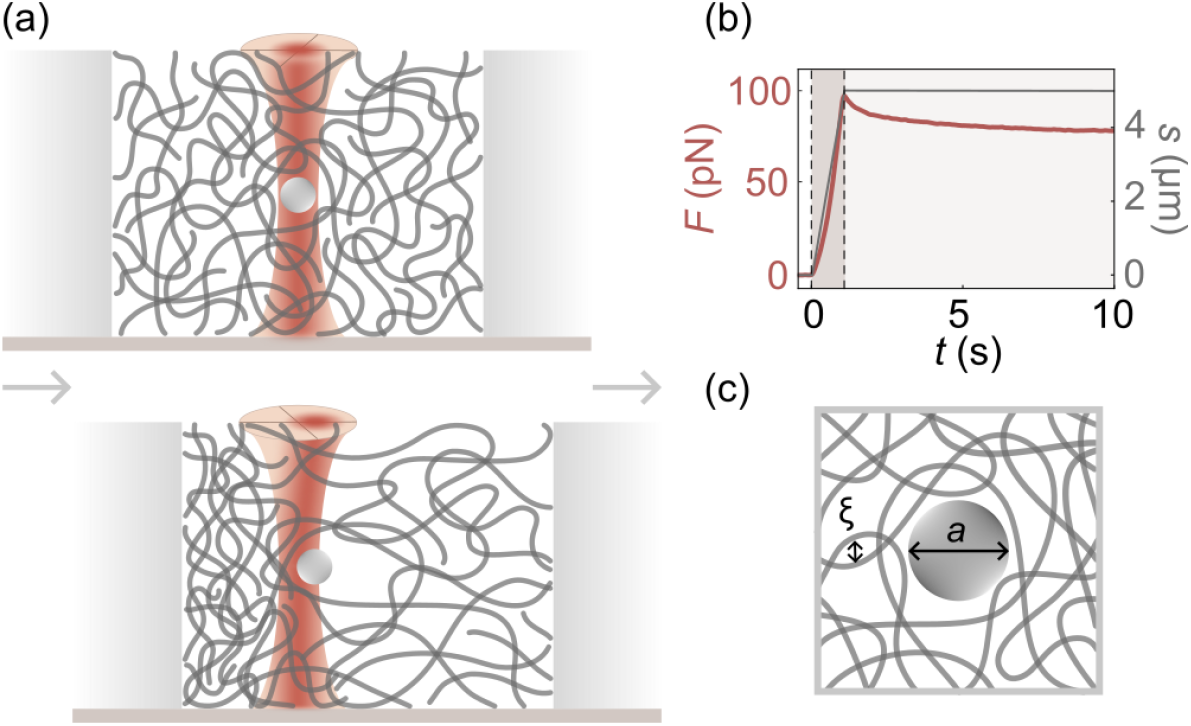
Experimental setup. (a) Schematic view of the AMR experiment for probing the filament networks using optical tweezers. Top: undisturbed bead; bottom: displaced bead relative to the network. (b) Example data from the AMR experiment showing the stage position *s* (gray) and the measured force *F* (red); the dark-shaded and light-shaded areas correspond to the extension and relaxation phases, respectively. (c) Sketch of the bead (diameter *a* = 2 *µ*m) within the network (mesh size *ξ* ~ 450 nm).

In Fig. 2(a) and (b) we show example data for vimentin and actin, respectively, for a maximum displacement of *s*_max_ = 5 *µ*m and a velocity of 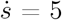 *µ*m/s. Each plot combines ten different measurements on ten different beads collected from the same sample chamber. We observe marked differences between the responses of the two networks. Pulling beads through vimentin networks requires forces that are considerably higher, by more than one order of magnitude, than for actin networks. Furthermore, the beads in the actin network relax back to zero force within a few seconds, a signature of “liquid-like”, viscous behavior. By contrast, the beads in the vimentin networks do not, not even after a relaxation time of 2 mins (SM, Fig. S2), indicating “solid-like”, elastic behavior. The data for vimentin networks are very smooth, while those for the actin networks are noisier due to the low force scales. Interestingly, we observe yield events, visible as “jumps” in the data during the extension phase in the actin data, especially at high displacements, see SM, Fig. S3.

**Figure 2:**
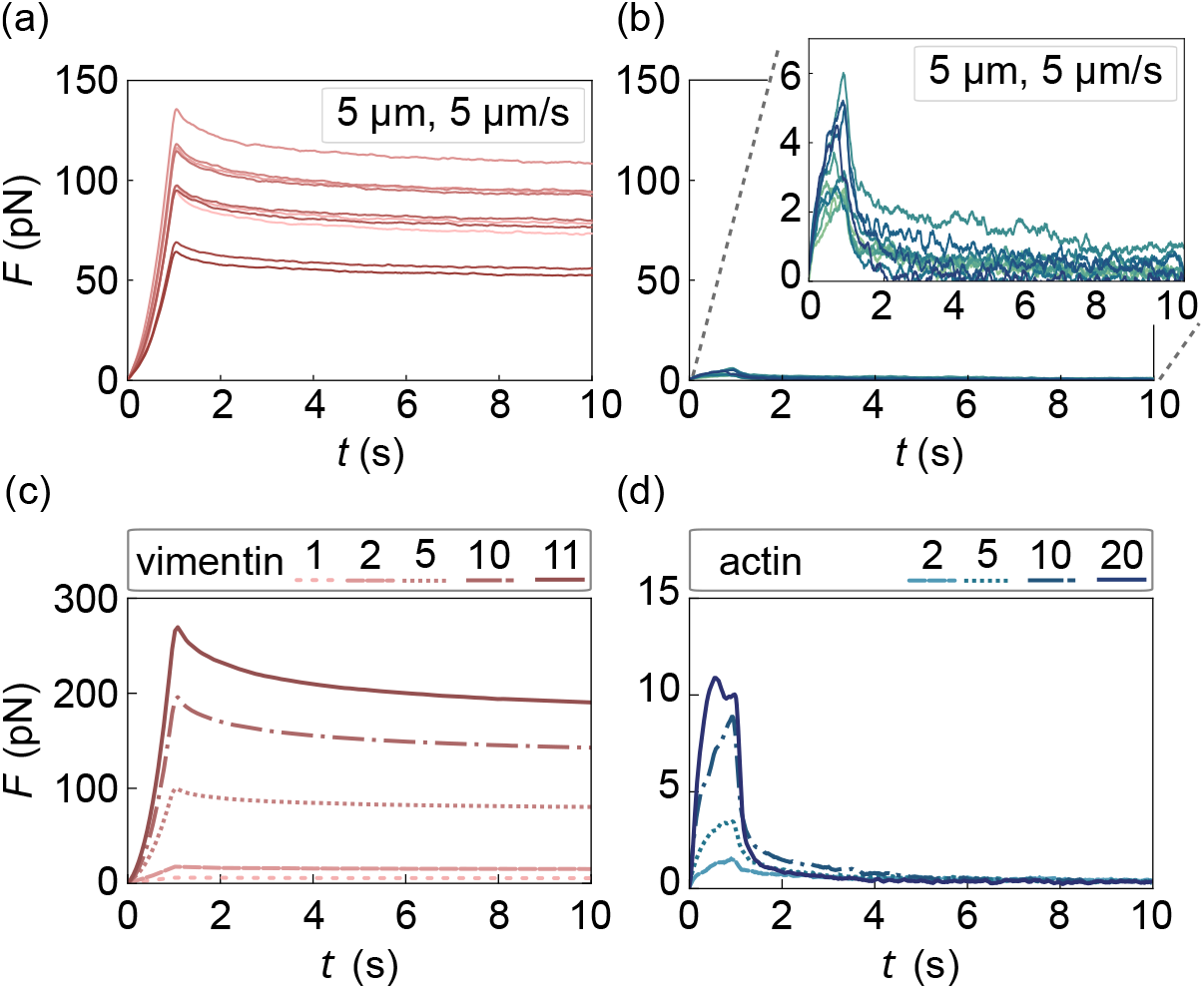
Force-time curves for (a,c) vimentin and (b,d) actin networks obtained from AMR experiments (protocol 1). (a,b) Individual measurements for a maximum displacement *s*_max_ of 5 *µ*m at a velocity 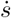 of 5 *µ*m/s. Each curve stems from a different bead within the same network. In (b) the inset shows zoomed versions of the same data for better visibility. (c,d) Average force-time curves for different displacements *s*_max_ and velocities 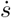 (see legend for color code and SM, Fig. S3 for the corresponding individual measurements).

Fig. 2(c) and (d) show average force-time curves for vimentin and actin, respectively, for increasing velocities and maximum displacements. As the vimentin networks are more strain-resistant than the actin networks, they require considerably higher forces for deformation. To remain within safe laser power limits we therefore restrict the maximum displacement for vimentin networks to 11 *µ*m, while for actin networks we go up to 20 *µ*m.

While Fig. 2 shows data for both the extension and relaxation phases, Fig. 3 concentrates on the extension phase, where the network is actively strained. To investigate whether the network response to strain is velocity dependent, we proceed as in previous studies of single vimentin filaments^7^ and keep the maximum distance constant while varying the pulling velocity [Figs. 3(a,b)]. Each curve is averaged across 10 different measurements. Note that the vimentin data [Fig. 3(a)] for a high velocity of 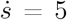 *µ*m/s end prematurely at less than the maximum distance of *s*_max_ = 10 *µ*m, because the beads escaped the traps, likely due to stiffening of the network. Indeed, for vimentin we observe a pulling speed (loading rate) dependence, indicating that when these networks are perturbed faster, they stiffen and require higher forces as compared to perturbations that occur more slowly. The data for 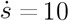 *µ*m/s are not included in this analysis as inconclusive behaviors were observed, possibly due to disruption of the networks structure at such high velocities.

**Figure 3:**
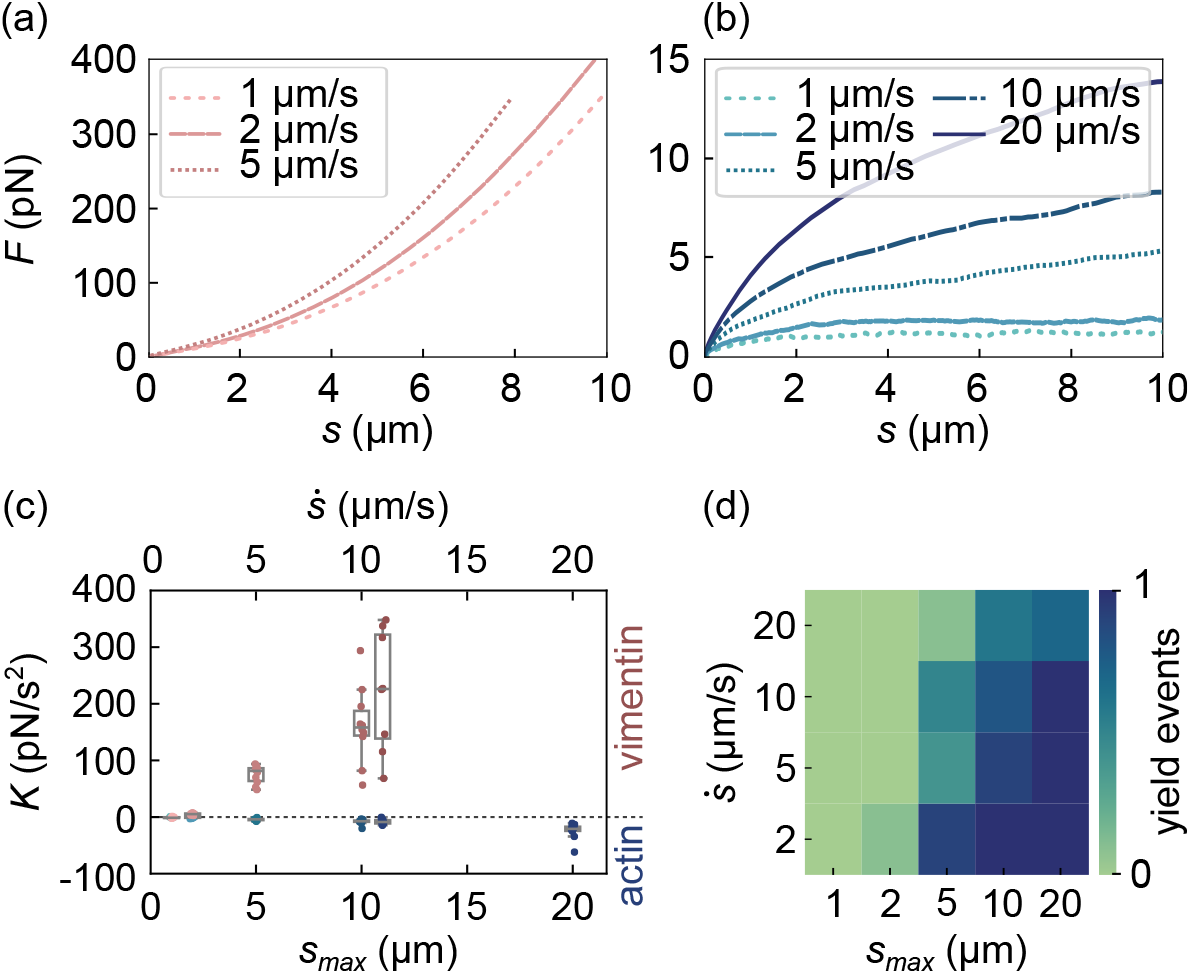
Comparison of AMR results for vimentin (red) and actin (blue) networks during the extension phase. (a, b) Force-displacement curves at different velocities (protocol 2), see legend for color code) for vimentin and actin, respectively. Each curve represents the average of ten individual curves. (c) Curvatures *K* of the force data shown in Fig. 2 and SM, Fig. S3 during the first second of the extension phase (protocol 1) plotted against the maximum displacement *s*_max_ (bottom axis) and velocity 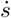 (top axis). The box plots show the quartiles and median. (d) Analysis of the actin data (protocol 2); the heat map shows the fraction of yield events at different maximum displacements and velocities; 0 (green) indicates no yield events (0), 1 (blue) indicates detected yield events in all cases (SM, Sec. IIBb)

Moving to the actin networks [blue, (b)], we also observe a loading rate dependence there, with a consistent trend across all investigated velocities between 1 and 20 *µ*m/s. For velocities of 1 and 2 *µ*m/s the average data curves reach a force plateau at about *s* = 2 *µ*m, with no further increase beyond and thus indicating that the network behaves as a viscous fluid. For larger velocities, such behavior is not observed, but instead the force increases with the displacement, indicating an elastic contribution to the mechanical behavior.

A striking difference between the vimentin and actin networks is that we consistently observe convex and concave force-displacement curves, respectively, during the extension phase (Fig. 2 and SM, Fig. S3). The convex curves for vimentin indicate strain-stiffening behavior, while the concave ones for actin indicate that the network acts predominantly like a fluid. In Fig. 3(c), to quantify these characteristics of the networks, we present the curvature *K* via the second derivative of the force-versus-time data from protocol 1. Clearly, at *s*_max_ of 1 *µ*m and 2 *µ*m, we observe no significant curvature, and between 2 *µ*m and 5 *µ*m, the curves start to deviate from the linear behavior: the vimentin data extend to high positive values, whereas the actin data show small negative values. For the larger distances and velocities the data scatter more, because the degree of heterogeneity in the networks becomes more evident.

A consistent feature of the vimentin data is the smooth increase in force with increasing displacement during the extension phase. In contrast, the actin data show frequent and sudden yield events, characterized by a peak in force, followed by a sudden drop and subsequent rise, as plotted in Fig. 3(d) as a heat map of 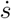 versus *s*_max_. The blue region represents the experimental parameters where such yield events contribute significantly to the stress relaxation: one sees that this is the case for large *s*_max_ and small 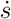. These observations can be rationalized by assuming that the network cannot relax stress on arbitrarily small timescales but requires some minimum relaxation time *τ*_0_ for this. Such a timescale for local plastic yielding is in fact a standard ingredient in elastoplastic models for the rheology of amorphous materials.^18–20^ Applying this to our experimental setting, yield events should typically be observed only when the extension phase is longer than *τ*_0_, i.e., when 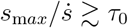, consistent with the trend seen in Fig. 3(d). From a linear representation of the data [SM, Fig. S4(a)] we find that the estimate *τ*_0_ ≈ 0.3 s gives a reasonable qualitative account of our experimental results; this timescale can also be estimated independently from relaxation phase data [SM, Fig. S4(b)]. Looking finally at the force levels where yield events occur for actin, we note that these are consistent with the forces where single actin filaments break when stretched using dual-trap optical tweezers (SM, Fig. S1).

To investigate how actin and vimentin networks release stress, we stop the stage movement after 1 s. Data for relaxation after an extension phase at a velocity of 1 to 11 *µ*m/s for vimentin and 2 to 20 *µ*m/s for actin networks are shown in Fig. 4 [(a) linear plot, (b) double-logarithmic plot; see color code in legend]. For faster initial pulling, the relaxation occurs faster. To confirm that the effect is strain rate – and not strain – dependent, we perform additional experiments, where the original displacement is 6 *µ*m for different velocities (protocol 2, SM, Fig. S5). We normalize *F* by the force *F*_0_ at the beginning of the relaxation phase to increase comparability of the individual curves. Vimentin networks do not release the stress completely, whereas actin networks fully relax. Moreover, the actin networks release the stress faster than the vimentin networks, i.e., the curves decay more rapidly.

**Figure 4:**
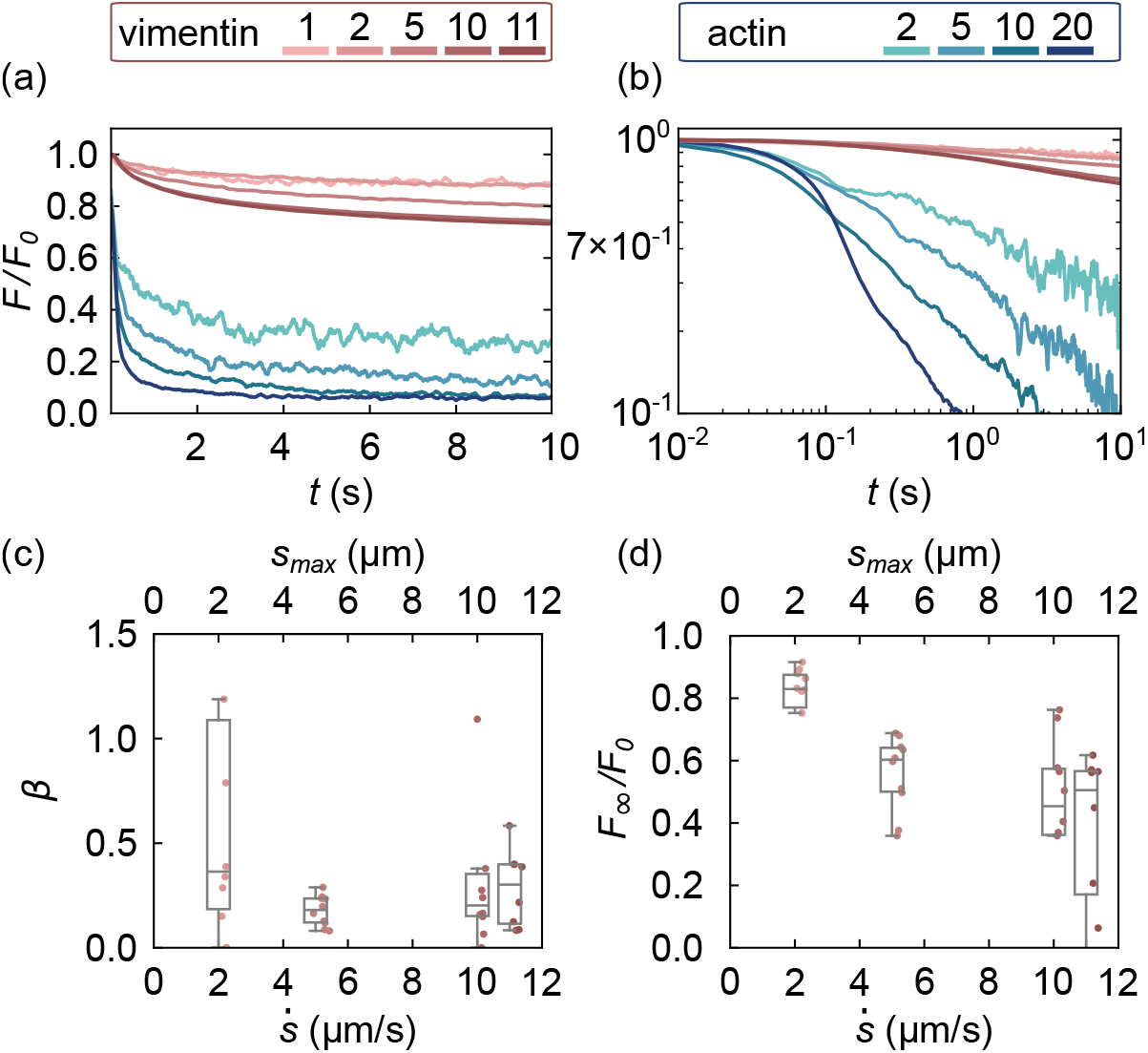
Comparison between AMR results of vimentin (red) and actin (blue) networks during the relaxation phase after a total extension time of *t* = 1 s (protocol 1). (a) Linear plot and (b) log-log plot of the relaxation data normalized by the initial force *F*_0_. Each curve represents the average of ten individual measurements. (c) Exponent *β* and (d) normalized force values at infinite time *F*_∞_*/F*_0_ for different maximum displacements *s*_max_ (top axis) and velocities 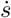 (bottom axis). The box plots show the quartiles and median. The circles represent individual data points.

Fig. 4(a) shows that for vimentin the curves of *F/F*_0_ corresponding to velocities of 1 and 2 *µ*m/s overlap; however, for higher velocities and maximum displacements the network releases the stress faster. This might indicate that the enhanced probability of local relaxation, i.e., breaking or rearranging of filaments, moves the material away from the linear rheology regime. Such behavior is not uncommon in soft biological matter systems. ^19,20^ We quantitatively analyze the data using a corresponding fit function:

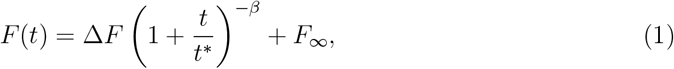

with Δ*F* = *F*_0_ − *F*_∞_. Here the plateau value *F*_∞_ accounts for the fact that the vimentin data do not generally relax to zero force, see double-logarithmic plot of the data in Fig. 4(b), and data for 60 s relaxation time in SM, Fig. S6. *F*_0_ is the force at the beginning of the relaxation phase and *t*^∗^ sets an overall time scale for the relaxation. Smaller values of *t*^∗^ therefore correspond to faster relaxation, as do larger values of the exponent *β >* 0 governing the power law part of the relaxation. Intuitively, the force relaxation described by Eq. (1) can be viewed as a superposition of an elastic, non-relaxing contribution of relative weight *F*_∞_*/F*_0_, and a set of Maxwell modes with a distribution of relaxation times *τ*. This distribution is peaked around *t*^∗^ and has a power law tail *τ*^−(*β*+1)^ at large *τ* (see SM, Sec. IIIB for details). Consistently with the trends discussed above, larger *β* thus corresponds to a relaxation time distribution that is less spread out towards values of *τ* above *t*^∗^, giving a faster decay of *F* (*t*).

By performing least-squared fits of Eq. 1 to the experimental data for *F* (*t*)*/F*_0_, we obtain *β* and *F*_∞_*/F*_0_ for the vimentin networks, probed at different experimental parameters, as shown in Fig. 4(c,d). Away from the region of small *s*_max_, where fit values for *β* have a wide spread due to increased noise levels, we observe an overall trend of *β* increasing with displacement or velocity. *F*_∞_*/F*_0_ indicates the fraction of the initial force that is not relaxed at long times and decreases with larger velocities and strains. Our data thus show that the network has a predominantly viscous rather than elastic response at higher velocity/higher strain, indicating it has a greater overall capacity to relax stress and to do so quickly.

The strain stiffening and softening in the vimentin and actin networks, respectively, is in good agreement with bulk shear rheology data.^2^ Our results also agree with previous AMR studies on actin concerning the strain rate dependence of reaching the viscous regime and stress softening after 0.01 s of the extension phase,^9^ although we used smaller beads (2 *µ*m versus 4.5 *µ*m) at a similar mesh size.^10^ Our estimated relaxation time on the order of 0.3 s is also in line with earlier work on entangled actin networks.^21^ This agreement sets the stage for the interpretation of the vimentin data. Similar experiments to ours have been conducted by Pinchiaroli et al.;^11^ however, in stark contrast to our results, the authors observe lower resisting forces by pure vimentin than by pure actin networks and, when mixing the networks, a long-time stress response that is governed by actin. Interestingly at low or high frequencies the authors observe more fluid- or solid-like behavior (*G*^*′′*^ *> G*^*′*^ or *G*^*′′*^ *< G*^*′*^), respectively, thus quite the opposite of what was found in in vitro vimentin networks^13^ and in cells.^22^ We believe that these differences arise from differing experimental procedures. Ref.^13^ shows that vimentin networks mature on a time scale of many hours, which is why in the present work we chose to assemble the vimentin networks for 22 hours, whereas in Ref.^11^ the assembly time was only 30 mins. Moreover, we assemble the networks around the beads within the measurement chamber, whereas in Ref. ^11^ they were first assembled and then flowed into the chamber, which might impose shear on the filaments and networks. We are aware that even after an assembly time of 22 h the networks continue to mature.^13^ Additionally, the energy input by the laser increases the temperature and thereby accelerates the maturation. When performing experiments with a relaxation phase of 15 s, and thus a total experiment time of about 1.5 h for all parameter sets, we observe no change as shown by repeating a measurement for the first parameter set at the very end of the experiment [SM, Fig. S7(a)]. When increasing the relaxation phase, and thereby the total experiment time to about 2.5 h, there is a stiffening observed [SM, Fig. S7(b)]). However, our conclusions remain unchanged. Comparing our data to the AMR study inside vimentin-free fibroblast cells compared to actin-free ghost cells,^12^ we find excellent agreement. The authors use a similar optical-tweezers approach as we do and move beads through the cytosol of the cells at constant velocity. The detected force range in the extension phase of our data aligns well with their experiments on ghost cells and relaxation is slower in ghost cells than in vim−/− cells. Additionally, they observe strain stiffening and softening in the vimentin and actin networks, respectively, just like we do.

In conclusion, we show that vimentin networks are mechanically distinct from actin networks. For high strains, where actin networks fluidize rapidly, vimentin networks show elastic behavior. The intriguing single filament mechanics of vimentin is also observed in the network scale, including the enormous extensibility, resistance to breaking and loading-rate dependence. In particular, vimentin networks are softer when they are deformed slowly, i.e., allowing for flexibility, and become stiffer when they are deformed fast, thus protecting the cells. This confirms the idea of a “safety belt” ^7,23–25^ for the cell and supports the notion that actin and vimentin have complementary roles in the cytoskeleton. Actin dynamically assembles and disassembles and rapidly rebuilds structures after damage. By contrast, vimentin networks – and possibly IF networks in general – are extensible and soft at slow velocities, but stiffen at high strains and at high velocities to ensure protection of the cell from damage.

## Acknowledgement

We thank Timo Betz and Bart Vos for helpful discussions, and Kamila Sabagh and Peter Luley for technical support. This work was funded by the Deutsche Forschungsgemeinschaft (DFG, German Research Foundation): Project-ID 449750155 - RTG 2756, projects A5 and A7 and by the European Union’s Horizon Europe program under the Marie Skłodowska-Curie Actions (MSCA), grant number 101148781. This research was conducted within the Max Planck School Matter to Life supported by the German Federal Ministry of Education and Research in collaboration with the Max Planck Society.

## Notes

### Competing Interest Statement

The authors have declared no competing interest.

