## Supplementary_Information for "Vimentin networks at high strains"

### Materials and methods

#### Bead passivation

Carboxylate-modified microspheres (2 $\mu$m-diameter, 2% (w/v), F8827, Life Technologies, Carlsbad, CA, USA) are passivated with $\alpha$-methoxy-$\omega$-amino polyethylene glycol ($CH_3O$˘PEG˘$NH_2$, 122000-2, Rapp Polymere, Tübingen, Germany) using 1-ethyl-3-(3-dimethylaminopropyl)carbodiimide (EDC; 22980, Thermo Fischer Scientific, Waltham, MA, USA) and N-hydroxysuccinimide (NHS; 24500, Thermo Fischer Scientific, Waltham, MA, USA) to form a covalent bond resistant to ion interference in the assembly buffer.[1] First, the beads are washed two times with 50 mM 2-(N-morpholino)ethanesulfonic acid (MES; 41320078-1, Bio World, Dublin, OH, USA) buffer at pH 6 by diluting 250 $\mu$L of bead solution in 750 $\mu$L of MES buffer, centrifuged at 9000 $\times$ g for 1 minute, and the pellet is resuspended in 1 mL MES buffer. Second, the carboxyl groups are activated through an EDC-NHS reaction. 20 $\mu$L of NHS solution in MES buffer at a concentration of 5.6 M and 20 $\mu$L of EDC solution in MES buffer at a concentration of 3 M are added to the bead solution. The mixture is incubated for 15 minutes at 37°C and 600 rpm on a shaker (ThermoMixer C, Eppendorf SE, Hamburg, Germany). Third, the sample is centrifuged at 9000 $\times$ g for 1 minute, the supernatant is discarded, and the pellet is resuspended in 1 mL MES buffer, vortexed and then sonicated for 15 seconds; this washing step is repeated twice. Fourth, after the second washing step, the pellet is resuspended in 500 $\mu$L borate buffer (0.05 M, pH 8.5) (bioPLUS, 40220048, BioWorld, Dublin, OH, USA), vortexed and sonicated for 15 seconds. For passivating the beads, 40 $\mu$L of coating solution ($CH_3O$˘PEG˘$NH_2$ in borate buffer at a concentration of 0.297 M) is added to the bead solution and incubated for 90–150 minutes at 37°C at 600 rpm followed by five centrifugation-resuspension cycles as explained in the washing step. Finally the bead pellet is resuspended in 250 $\mu$L MES buffer and stored at 4°C.

#### Protein preparation

Human vimentin mutant C328A is purified from inclusion bodies as described before.[2,3] For the microrheology experiments vimentin is reconstituted by dialysis in a stepwise manner from 8 M urea (8 M, 6 M, 4 M, 2 M, 1 M) into 2 mM phosphate buffer (PB), pH 7.5 to form tetramers. The protein concentration is adjusted using 2 mM PB and verified by UV absorption spectroscopy at 280 nm (Nanodrop ND-1000, ThermoScientific Technologies, Inc., Wilmington, DE, USA). For a network with a theoretical mesh size of 450 nm, a vimentin concentration of 1 g/L is required.[4,5] For the experiment, 5 $\mu$L of passivated beads are centrifuged at 9000 $\times$ g for 1 minute and resuspended in 400 $\mu$L of 2 mM PB at pH 7.5. We assemble vimentin networks in a custom microrheology chamber for 22 h at room temperature at a final concentration of 100 mM KCl by mixing the protein solution at 1.28 g/L with 6$\times$ assembly buffer (600 mM KCl in 2 mM PB at pH 7.5) and the passivated beads at a ratio of 19:4:1.

Actin is purified from muscle tissue as describe before.[6,7] G-actin at a concentration of 0.78 g/L, verified by UV absorption spectroscopy at 280 nm, is incubated on ice for 2 h in G-buffer (5 mM TRIS, pH 8.0, 0.2 mM ATP, 0.2 mM $CaCl_2$, 0.5 mM DTT) to depolymerize any aggregates and to reduce the number of nuclei. For the active microrheology (AMR) experiments, actin polymerization is performed as follows; F-buffer (10 mM TRIS, pH 7.5, 1 mM ATP, 2 mM $MgCl_2$, 50 mM KCl) is supplemented with passivated beads: 5 $\mu$L of passivated beads in MES buffer are centrifuged at 9000 $\times$ g for 1 minute and resuspended into 400 $\mu$L of 2$\times$ F-buffer. This suspension is then further diluted in 2$\times$ F-buffer at a ratio of 1:9. The assembly is initiated in a custom microrheology chamber for 3 h at room temperature. The final concentration of 0.39 mg/mL is achieved by mixing G-actin and 2$\times$ F-buffer with passivated beads at a ratio of 1:1 and provides the same mesh size of 450 nm as vimentin at 1 g/L.

For stretching single actin filaments, unlabeled G-actin, 3% fluorescently labeled actin and 30% biotin-labeled actin are mixed in G-buffer at a total concentration of 0.2 mg/mL

(Cytoskeleton, Denver, CO, USA). The mixed solution is placed on ice for one to two hours to depolymerize actin oligomers that have formed during storage. Polymerization is initiated by adding 10× F-buffer (volume ratio actin in G-buffer : 10× F-buffer = 9 : 1) and the mixture is incubated at room temperature for one hour. The resulting filaments are diluted to 70 nM in F-buffer and stabilized by adding phalloidin in abundance (Fisher Scientific, Schwerte, Germany). Filament formation is confirmed prior to the measurements using epi-fluorescence microscopy.

#### Microrheology chamber preparation

1 mm microscopy slides ($24 \times 60$ mm$^2$; VWR International, Radnor, PA, USA) and no. 1.5 coverslips ($18 \times 18$ mm$^2$; VWR International) are sterilized in isopropanol and dried with nitrogen. Double-sided tape (tesafix 4959, Tesa, Norderstedt, Germany) with 100 to 120 $\mu$m thickness is cut into a rectangle-shaped frame ($9 \times 8$ mm$^2$ inner frame size) and placed on the sterilized microscopy slide. 6.5 $\mu$L of either actin or vimentin solution is added in the center of the frame. A hydrophobic border is drawn on the sterilized coverslip using a 2 mm PAP pen (Sigma-Aldrich, St. Louis, MO, USA) to additionally seal the chamber. The chamber is then closed with the coverslip and placed on a custom-built rotor to prevent settling of the beads on the glass slides during incubation.

#### Active microrheology

We perform microrheology experiments using optical tweezers (C-Trap, Lumicks, Amsterdam, The Netherlands). The data are collected using a water immersion objective (60×, NA 1.2). To be able to quantify the force values by optical tweezers, we determine the conversion factor from the voltage recorded on the detector to the bead displacement in the trap and the trap stiffness in water from the power spectral density (PSD) of the thermal motion of the beads (passive microrheology (PMR) measurement). Ten data sets are recorded and averaged. Embedded beads in the networks are used as probe particles. For the measurements we choose beads that are located in the center of the chamber, about 30 $\mu$m away from each glass surface. A PMR experiment is conducted for 30 seconds before each AMR experiment to record the thermal fluctuations of the particle at the respective laser power and thereby ensure that the network is properly assembled.

For the AMR experiments, we capture a bead, keep it in place and move the stage (and thereby the chamber with the network) to the specified maximum displacement $s_{\max}$ at the specified speed $\dot{s}$. We use laser powers of 133-260 mW to ensure that the networks are not destroyed by heating. To observe the network's relaxation after mechanical straining, the stage movement is stopped, allowing the probe particle to relax within the trap while the force is recorded.

#### Stretching of single actin filaments

We stretch single actin filaments using a dual-trap optical tweezers system combined with confocal fluorescence microscopy and a microfluidic device (C-Trap, Edge 400, Lumicks). The experiments are performed in a three-inlet microfluidic flow cell. Channel 1 contains 4.5 $\mu$m streptavidin-coated polystyrene beads (Lumicks), diluted in G-buffer (1:200) to allow for covalent binding to the actin filaments via streptavidin-biotin interaction. Channel 2 contains the measuring buffer (2 mM PB (pH 7.5), 100 mM KCl, 1 mM $MgCl_2$, 1 mM DTT, and 0.2 mM ATP). Channel 3 contains polymerized actin filaments diluted 1:100 in assembly buffer and supplemented with an oxygen scavenger system (0.008 mg/mL catalase, 0.04 mg/mL glucose oxidase, 1.2 mg/mL glucose, 20 mM DTT, 5 mM TRIS (pH 8.0), 0.2 mM Na-ATP, 0.2 mM $CaCl_2$). To start a measurement, two beads are captured with the optical tweezers in channel 1. The trap stiffness is calibrated by analyzing the PSD of the thermal fluctuations in channel 2. Actin filaments are attached to the beads in channel 3, and confocal microscopy is used to verify that only a single actin filament is tethered between the beads. The traps with the actin filament are moved back to channel 2, and the flow is stopped. The filaments are stretched at a loading rate of $0.27 \pm 0.03\,\mu$m/s by moving one of the traps until they rupture.

### Data analysis

All data analysis is carried out with the self-written python codes.

#### Single stretched actin filaments

Force-distance curves are obtained by combining force data collected at 78 kHz with distance data derived from bead positions recorded at 4 Hz using bright field imaging. The force data are downsampled to 10 Hz using the `downsampled_by()` function from the Lumicks Pylake package.[8] To match the temporal resolution of the downsampled force data, the bead position-derived distances are linearly interpolated using `numpy.linspace`. From these synchronized force-distance data, force-strain curves are calculated at 10 Hz for each filament. The strain is defined as

$$\epsilon = \frac{L - L_0}{L_0}, \tag{1}$$

where $L$ is the measured extended filament length, and $L_0$ is the filament length at 1 pN.

#### Active microrheology

The displacement of the bead with respect to the trap center is measured by back-focal-plane interferometry with a quadrant position detector (QPD). Using the conversion factor and the trap stiffness, we apply Hooke's law to determine the force on the bead.

**Curvature in the extension phase** To estimate the curvature in the extension phase, we fit the following second-order polynomial to the first second of force-time data:

$$F(t) = K \cdot t^2 + m \cdot t + c \tag{2}$$

using a least-squares optimization. This yields the parameter $K$, which indicates the strength and direction (concave or convex) of the curvature.

Alternatively, one can define a dimensionless curvature $\tilde{K}$ as

$$\tilde{K} = \frac{K t_0^2}{\Delta F}, \tag{3}$$

which measures the curvature independently of the total force increase $\Delta F = F(t_0) - F(0)$ of the extension phase of duration $t_0$. We show the estimated values for $\tilde{K}$ for actin and vimentin in Fig. 8. Vimentin shows an increase in dimensionless (convex) curvature up to a displacement of 5 $\mu$m after which it plateaus. Actin shows consistently negative curvature (concave), without a trend that is perceivable within the noise range.

**Yield events in actin data** Ten force-time curves for each velocity (2, 5, 10, and 20 $\mu$m/s) and a total displacement of 20 $\mu$m are analyzed individually. "Yield events" are defined as negative slopes after a local maximum. Once a yield event occurs at a given displacement, it will also be counted as such at any larger displacement $s_{\max}$. Thus, Fig. 3(d) in the main text provides a quantitative representation of the probability of yield events to occur during the active movement. In other words, the color scale in the heat map in Fig. 3(d) in the main text corresponds to fractions of yield events between 0 (no yield events) and 1 (yield events for all individual curves within the averaged ensemble).

**Relaxation phase in vimentin data** All data are collected at a sampling frequency of 78 kHz. To divide the extension and relaxation phases, we use the time point where the nanostage changes its moving direction. Using least squares fitting of Eq. (1) in the main text to the individual data curves, we obtain the four free fitting parameters: $\beta$, $t^*$, $\Delta F$, and $F_\infty$. As an alternative, to check whether our data show significant signs of a finite rest-force (plateau), we consider a model with fixed $F_\infty = 0$ as well, and fit it to our experimental data. We measure the goodness-of-fit for both models using a $\chi^2$-test and find the reduced $\chi^2$-value to be 1.44 times larger in the case of the simplified model without $F_\infty$. This indicates larger discrepancies between the model and the data. Therefore, we use the more complex model, which better explains the observed force curves.

### Theory

#### Relaxation time in actin data

As discussed in the main text, our data on the occurrence of yield events in actin networks during the extension phase suggest that there is a typical minimal timescale $\tau_0$, or correspondingly a maximal rate $1/\tau_0$, for yield events to take place. Yield events are then only expected in experiments where the extension phase takes longer than $\tau_0$, i.e. where $\dot{s} < s_{\mathrm{max}}/\tau_0$. The data in Fig. 4(a) are broadly consistent with this, with $\tau_0$ of order 0.3 s.

We now show that this timescale can also be extracted from the relaxation phase data for actin, presented in blue in Fig. 4(a,b) in the main text. We focus on the larger displacements, where a crossover from exponential to power law relaxation is visible. The power law tail is reminiscent of soft glassy materials,[9,10] suggesting a fit to predictions of the soft glassy rheology (SGR) model.[11] In linear response the force after the displacement ramp in the extension phase is $F(t) \propto \dot{s} \int_{-t_0}^{0} dt'\, G(t-t')$, where we have chosen the time origin so that the extension phase runs from $t = -t_0$ to $t = 0$. We approximately account for nonlinearities from large displacements by using for the $G(t)$ the nonlinear SGR stress relaxation function for large step strains. Implementing a suggestion in Ref.[12] that the model should be modified to ensure that yield rates can be no larger than $1/\tau_0$, we find:[13]

$$G(t) = (1-w)e^{-t/\tau_0} + w\, G^{\mathrm{SGR}}(t/\tau_0) \tag{4}$$

where $G^{\mathrm{SGR}}$ is the linear stress relaxation function of the vanilla model and the weight $w$ decreases with strain amplitude, making the exponential part of the relaxation dominant for larger strains. By a least squares fit of the experimental data for $F/F_0$ [Fig. 4(a,b), blue] for a larger displacement ($t_0 = 1\,\mathrm{s}$, $\dot{s} = 10\,\mu\mathrm{m/s}$, $s_{\mathrm{max}} = 10\,\mu\mathrm{m}$) we then find $\tau_0 = 0.29\,\mathrm{s}$, consistently with the timescale for yield events extracted from the extension phase data.

#### Role of the exponent $\beta$

Power law linear rheology can be used to model materials with a broad distribution of relaxation times. A power law (shear stress) relaxation function $G(t) = (1 + t/t^*)^{-\beta}$ as used in Eq. (1) in the main text can be written as $G(t) = \int_0^\infty d\tau\, P(\tau) e^{-t/\tau}$ with a distribution $P(\tau)$ of relaxation times that is peaked around $t^*$ and has a power tail $\sim \tau^{-(\beta+1)}$ for large $\tau$. This form makes sense for arbitrary $\beta > 0$, and approaches an exponential (Maxwell) form $G(t) = e^{-\beta t/t^*}$ for large $\beta$.

The corresponding frequency dependent shear moduli $G^*(\omega)$, given by the Fourier transform of $G(t)$ up to a factor of $i\omega$, contain a low-frequency term $\sim (i\omega)^\beta$ with the same exponent $\beta$. One has to be careful not to conclude erroneously, however, that $\beta = 1$ corresponds to purely viscous relaxation: at $\beta = 1$ there are additional logarithmic factors $\ln(\omega)$ in $G^*(\omega)$, which arise from the competition of $(i\omega)^\beta$ with additional non-singular contributions $\sim i\omega$, $(i\omega)^2$ etc.

(13) Mendozza, R.; Sollich, P. in preparation.

### Supplementary figures

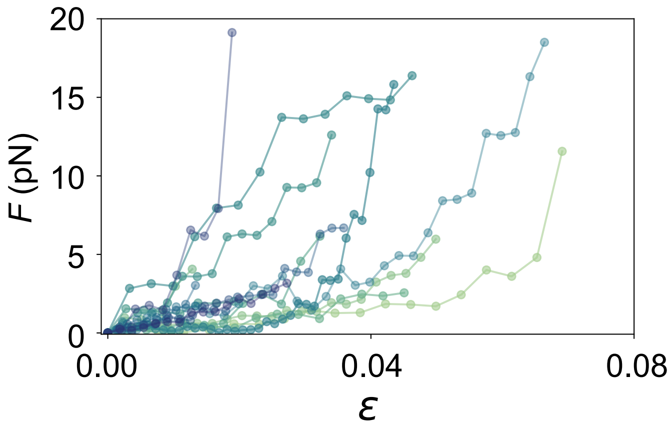


Figure 1: Force-strain plot for 16 individual actin filaments stretched at a loading rate of $0.27 \pm 0.03\,\mu$m/s using optical tweezers. The data points are connected as a guide to the eye. The filaments break at low forces on the order of a few pN and at low strains.

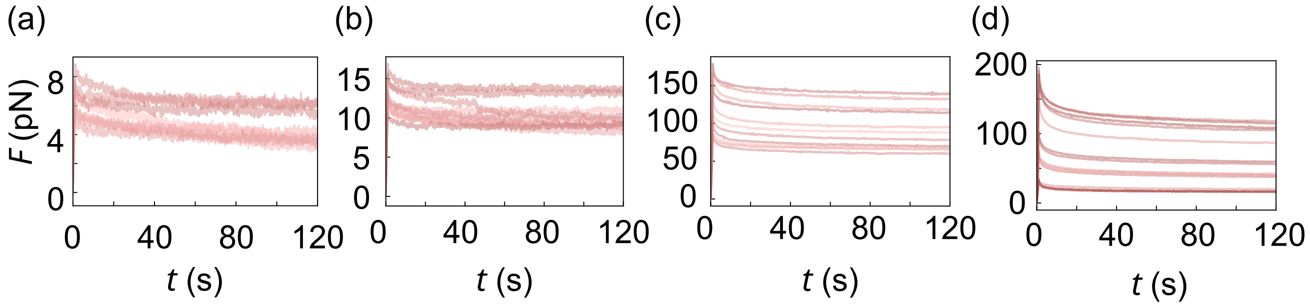


Figure 2: Force-time plots for vimentin networks at extended relaxation times of 120 s at different velocities $\dot{s}$ and displacements $s_{\max}$, at 1 s total extension time (protocol 1): (a) 1 $\mu$m, 1 $\mu$m/s, (b) 2 $\mu$m, 2 $\mu$m/s, (c) 5 $\mu$m, 5 $\mu$m/s, (d) 10 $\mu$m, 10 $\mu$m/s. Even at such long relaxation times, the data do not decay to zero force.

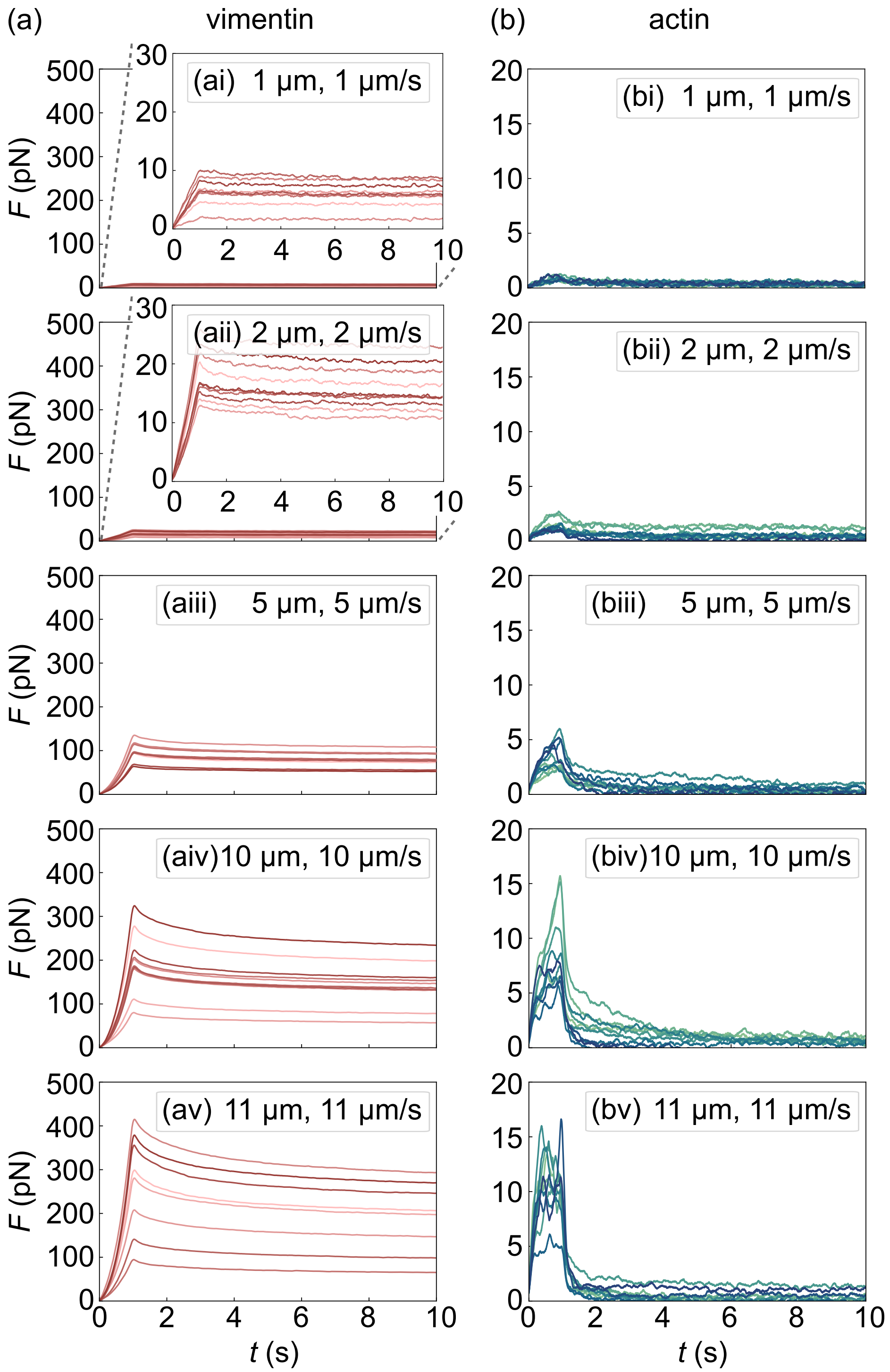


Figure 3: Force-time plots for (a) vimentin and (b) actin networks obtained from AMR experiments (protocol 1) for different velocities $\dot{s}$ and displacements $s_{\max}$ (as indicated in the individual panels), at 1 s total moving time. For each protein, all data are plotted on the same force scale. Note the different force scales in (a) and (b). In (ai,aii) the insets show zoomed versions of the same data for better visibility. Each sub-panel displays ten separate curves. The data clearly show a high degree of heterogeneity, which likely arises from the intrinsic heterogeneity of the reconstituted networks. Despite this heterogeneity, however, the overall force magnitude consistently increases with increasing speed and displacement for both network types

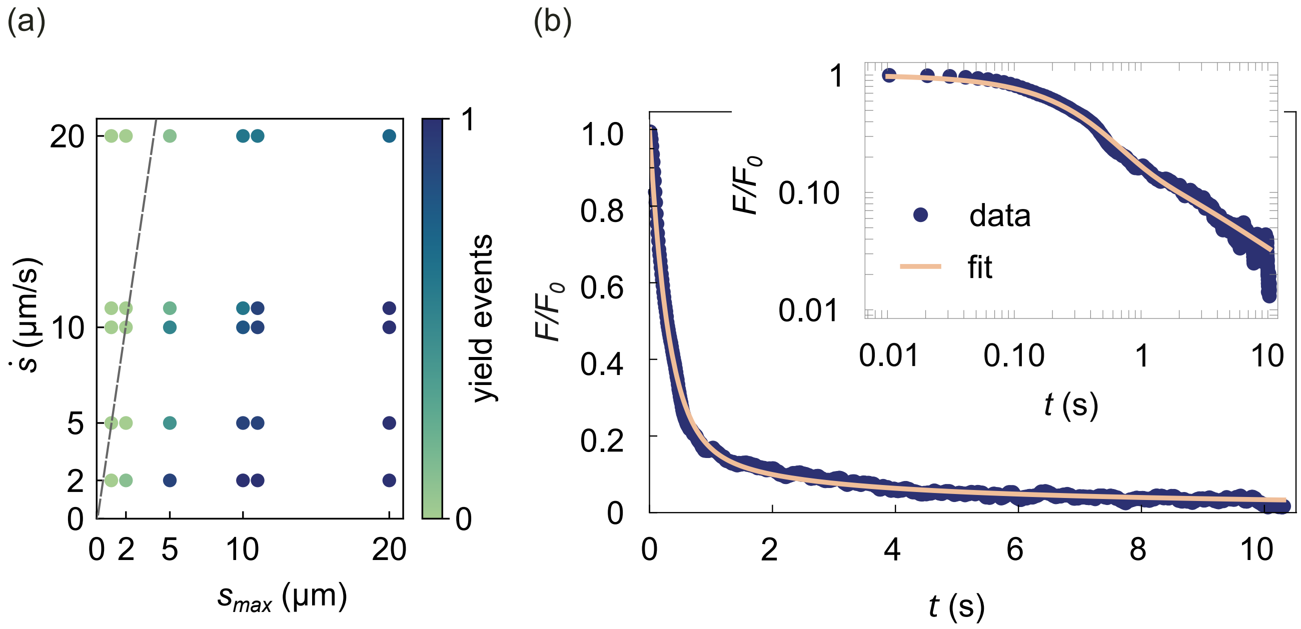


Figure 4: (a) Linear representation of the data shown in Fig. 3(d) in the main text. The dashed line corresponds to $\dot{s} = s_{\max}/\tau_0$, where $\tau_0 = 0.29$ s is obtained from the fit shown in panel b. One expects the probability for yield events to be significant only below this line, which is qualitatively consistent with the experimental data. (b) Main plot: Fit of the modified SGR model from Eq. (4) to actin force relaxation data subsequent to a loading phase with $\dot{s} = 10$ $\mu$m/s. Time $t = 0$ indicates the beginning of the relaxation phase, where $F = F_0$. Inset: log-log representation of the same data, illustrating that the model reproduces the crossover from exponential to power law relaxation.

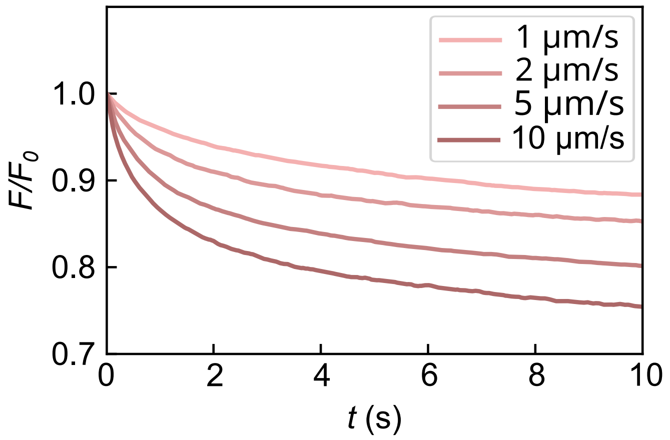


Figure 5: Velocity dependence of the relaxation phase in vimentin networks; shown for experiments with the same maximum displacement of 6 $\mu$m at different velocities during the extension part (protocol 2). Each curve represents the average of ten individual curves. These data show that the network relaxes faster after it has been deformed faster.

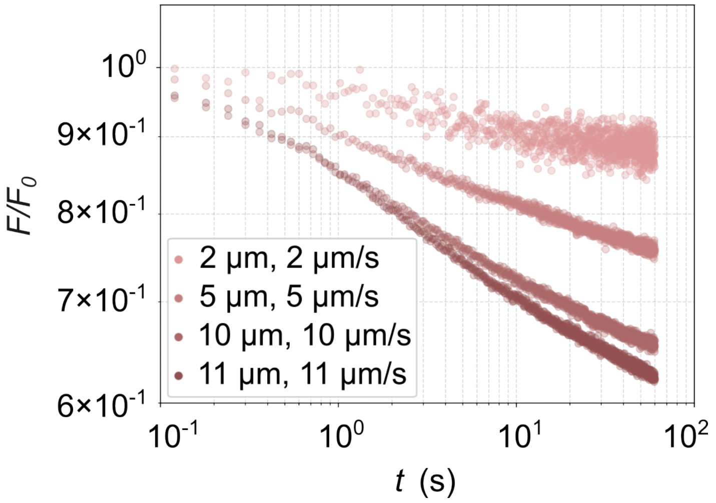


Figure 6: AMR experiment on vimentin networks. Data from Fig. 4(b) in the main text plotted for an extended relaxation time of 60 s. Each curve represents the average of ten individual measurements. For the small displacement (2 $\mu$m) and velocity (2 $\mu$m/s) the data reach a plateau at 1 min relaxation time, for the other parameter sets, even longer relaxation times are required to develop the plateau.

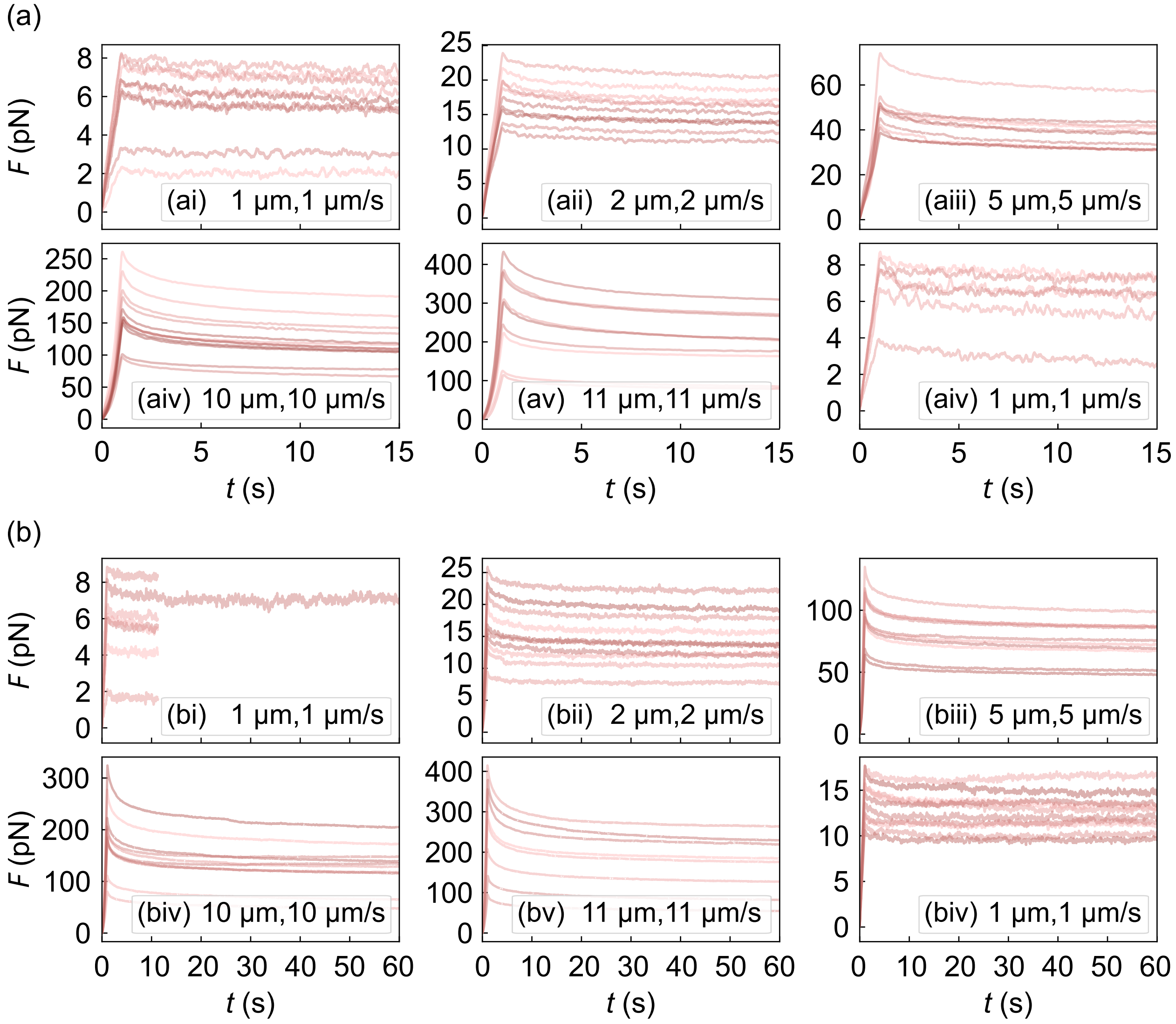


Figure 7: Series of Force-time plots for vimentin using different velocities $\dot{s}$ and displacements $s_{\max}$ (as indicated in the individual panels); in each case, the parameters in the last measurement (aiv, biv) correspond to the first one (ai, bi). This last measurement is performed to control for changes in the network due to further maturation during the experiment time. (a) Relaxation phase of 15 s, total experiment time approximately 1.5 h. No difference between the first and the last measurement is observed. (b) Relaxation phase of 60 s (note that in (bi) we did not record the data for this longer time), total experiment time approximately 2.5 h. The networks mature further during the measurements, due to the longer total time and energy input by the laser beam, and therefore stiffen; the forces detected in the last measurement are about two times as high as in the first measurement, albeit still lower than for the second parameter set (bii).

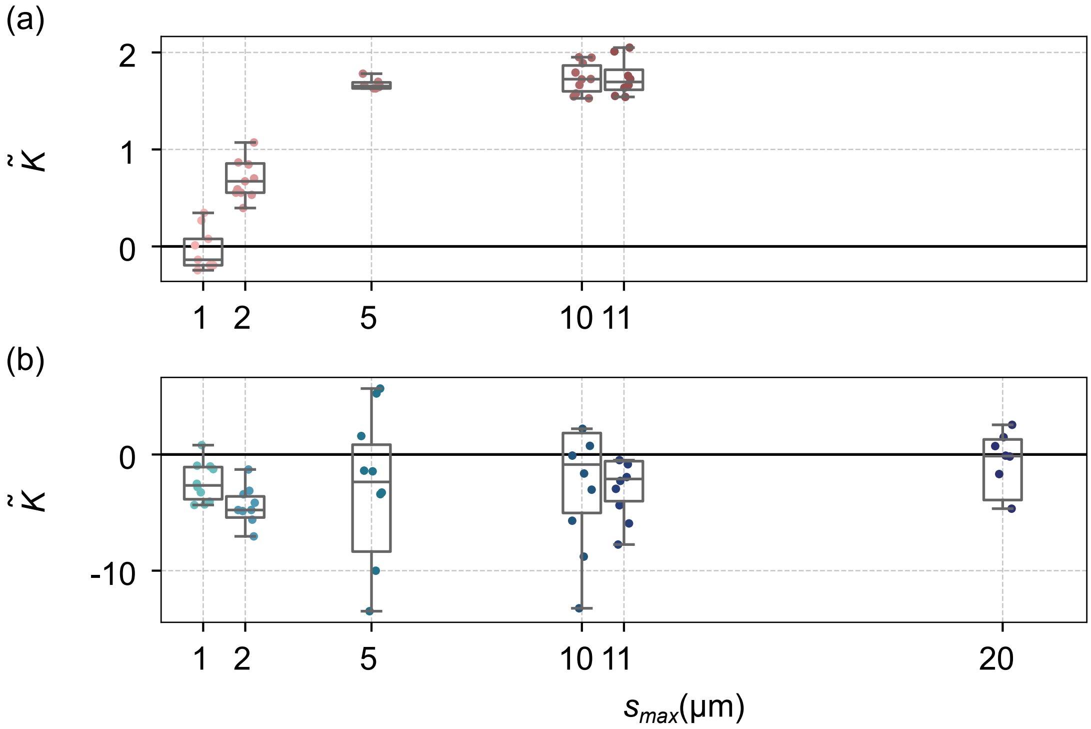


Figure 8: Dimensionless curvatures $\tilde{K}$ of the force data of the extension phase plotted against the maximum displacement $s_{\max}$, (a) vimentin, (b) actin. The datapoints are scattered over the width of the box for visibility.
